# Species-wide quantitative transcriptomes and proteomes reveal distinct genetic control of gene expression variation in yeast

**DOI:** 10.1101/2023.09.18.558197

**Authors:** E. Teyssonnière, P. Trébulle, J. Muenzner, V. Loegler, D. Ludwig, F. Amari, M. Mülleder, A. Friedrich, J. Hou, M. Ralser, J. Schacherer

## Abstract

Gene expression varies between individuals and corresponds to a key step linking genotypes to phenotypes. However, our knowledge regarding the species-wide genetic control of protein abundance, including its dependency on transcript levels, is very limited. Here, we have determined quantitative proteomes of a large population of 942 diverse natural Saccharomyces cerevisiae yeast isolates. We found that mRNA and protein abundances are weakly correlated at the population gene level. While the protein co-expression network recapitulates major biological functions, differential expression patterns reveal proteomic signatures related to specific populations. Comprehensive genetic association analyses highlight that genetic variants associated with variation in protein (pQTL) and transcript (eQTL) levels poorly overlap (3.6%). Our results demonstrate that transcriptome and proteome are governed by distinct genetic bases, likely explained by protein turnover. It also highlights the importance of integrating these different levels of gene expression to better understand the genotype-phenotype relationship.

**Highlights:**

- At the level of individual genes, the abundance of transcripts and proteins is weakly correlated within a species (*ρ* = 0.165).
- While the proteome is not imprinted by population structure, co-expression patterns recapitulate the cellular functional landscape
- Wild populations exhibit a higher abundance of respiration-related proteins compared to domesticated populations
- Loci that influence protein abundance differ from those that impact transcript levels, likely because of protein turnover

## Introduction

Understanding the genetic basis of phenotypic variation in natural populations is one of the main goals of modern biology. Gene expression differs among individuals and is known to be a main determinant of phenotypic variation^1, 2^. In humans, the onset and development of numerous diseases have been linked to abnormal regulation of gene expression^3^. It is therefore essential to understand how genomic information is expressed through the different layers of gene regulation (*i.e.,* transcriptomes and proteomes). Over the past decades, the development of methods for high-throughput quantification of mRNA and protein abundance has made it possible to explore both the proteome and the transcriptome on a larger scale^4, 5^. These approaches facilitated the detection of numerous genetic loci (quantitative trait loci, QTL) affecting either transcript (eQTL) or protein (pQTL) levels^6–11^. However, the relationship between transcript and protein levels remains debated and poorly understood at the population level^12^.

The transcript-protein correlation provides a first global view of the dependency of the two gene expression layers. Two types of mRNA-protein correlation can be determined, across- and within-gene, reflecting very different dynamics^12–14^. The across-gene correlation analysis focuses on the overall correlation of a large set of genes coming from the same sample under a given condition to find out how well the absolute abundances of mRNAs and proteins are correlated. This correlation has been widely investigated in several species, such as human^15–21^, rats and mice^22–25^, flies^26^, plants^27^ or yeast^28–30^. Across-gene correlations are consistently high and range from 0.4 to 0.8, suggesting that the absolute number of transcripts and proteins are globally correlated. Therefore, very abundant transcripts generally lead to very abundant proteins and vice versa.

However, the relationship between the transcript and protein abundance at the population level is explored via their variation across samples (*e.g.*, individuals, tissues, or cell lines). Within-gene correlation analysis gives a view on how the protein level of each gene tracks its mRNA level in a population. Different studies have investigated this within-gene correlation in different contexts and organisms, but they often show divergent results. Several surveys of tumors, normal human tissues, as well as pluripotent stem cells have highlighted this discrepancy in estimates with median within-gene correlation coefficients ranging from 0.14 to 0.59^15, 19, 21, 22, 31–39^. Similarly, the overlap of the detected loci influencing mRNA (eQTL) and protein (pQTL) abundance greatly differed across the datasets. It ranges from a very weak overlap of 5.5% in a study on 97 inbred and recombinant mice to nearly 35% in human (n = 62) and mice (n = 192)^6, 15, 40^.

Part of the diverging results might have been driven by technical limitations. For instance, it has been shown that by selecting the most representative peptides in prior proteomic methods, the overall correlation of global transcript and mRNA abundance improves significantly^37, 41^. A key difference is also whether the goal of the survey is to correlate absolute number of transcripts and proteins, or relative changes in protein or mRNA levels, which differ between samples. While the absolute number of transcripts and proteins spans several orders of magnitude, the relative expression differences of any individual protein across samples varies within a much narrower range^30, 42^. Finally, a main limitation of these studies is that the sample size is much lower than the dimensionality of the problem.

To determine to which extent differences in relative changes in mRNA and protein levels are correlated and the genetic origins of their abundance variation are shared, a large-scale population survey exploring these two facets in a quantitative way was therefore necessary. Here, we took advantage of the 1,011 yeast *Saccharomyces cerevisiae* population we genome-sequenced and for which we have a species-level understanding of the natural genetic and phenotypic diversity^43^. In order to be fully able to compare and analyze at unprecedented detail the relationship between these two layers of gene regulation, we therefore generated 942 quantitative proteomes in which cells were also cultured in synthetic complete medium supplemented with amino acids using high-throughput mass-spectrometry. We found that protein levels are molecular traits that exhibit considerable variation between individuals and specific signatures related to certain subpopulations. This large available population also makes it possible to generate a detailed map of loci involved in the variation of protein abundance (pQTL) at the species level, via genome-wide association studies (GWAS). Interestingly, local pQTL are less frequent than distant ones (8% of the total set of pQTL) but they have a higher impact on their respective traits. Integration of proteomic and transcriptomic datasets acquired in parallel under similar conditions allowed comparison of accurate quantification of the mRNA and protein abundance of 629 genes across 889 natural isolates^44^. Based on these unique datasets, we clearly demonstrated that the degree of within-gene correlation between protein and mRNA abundance is very low (*ρ* = 0.165). Consistently, we found that the genetic variants influencing protein and mRNA abundance are very dissimilar. Our study highlights that population-scale proteomes are essential and add a new dimension to the characterization of the genotype-phenotype relationship when integrated with genomic and transcriptomic information.

## Results

### Quantitative proteomes of a large collection of natural isolates

We generated a quantitative proteomic dataset for strains of the 1,011 strains collection^43^ from cells cultivated in synthetic complete medium with amino acids in order to match the growth medium used for RNA sequencing^44^ (Figure 1A).We had previously acquired a proteome dataset of the 1,011 strains collection, measured with microflow chromatography and SWATH MS^45^. For the acquisition of this new dataset we used a proteomic method that allows for an even higher throughput, using analytical flowrate chromatography and Scanning-SWATH MS with a 3 min gradient^46^. After cultivation of the yeast isolates in 96 well plates, proteins were extracted, and subjected to reduction, alkylation, and trypsination in a semi-automated workflow using liquid handling robotics^47^. Peptide preparations were separated using a 3-minute high-flow rate (800 µl/min) chromatographic gradient using an Infinity II chromatographic system (Agilent Technologies), coupled to a 6600 Triple TOF instrument (Sciex). Data was recorded using Scanning SWATH acquisition^46^ and the raw data was processed using the DIA-NN software (version 1.8), which was specifically developed for large scale proteomic exploration^48^. We applied several quality filters where poor-quality samples were removed from the analysis, and we excluded peptides that were not detected in more than 80% of the samples (see Methods). The generated dataset hence encompasses protein abundance quantification for 630 proteins among 942 isolates (Table S1 and Table S2). This dataset therefore covers the overall genetic diversity of the species and captures the subpopulations that were defined as part of the 1,011 yeast genomes project, including both domesticated and wild clades^43^ (Figure S1A). We combined the proteomic dataset with transcriptomic data obtained from the 1,011 strains collection^44^, which gave access to the quantified expression of both levels for 629 genes across 889 isolates (Figure 1B, Table S1). To be able to properly compare these two datasets, we normalized them with quantile normalization after imputing the missing values using the KNN method (Table S3, Figure S1B-C).

**Figure 1.**
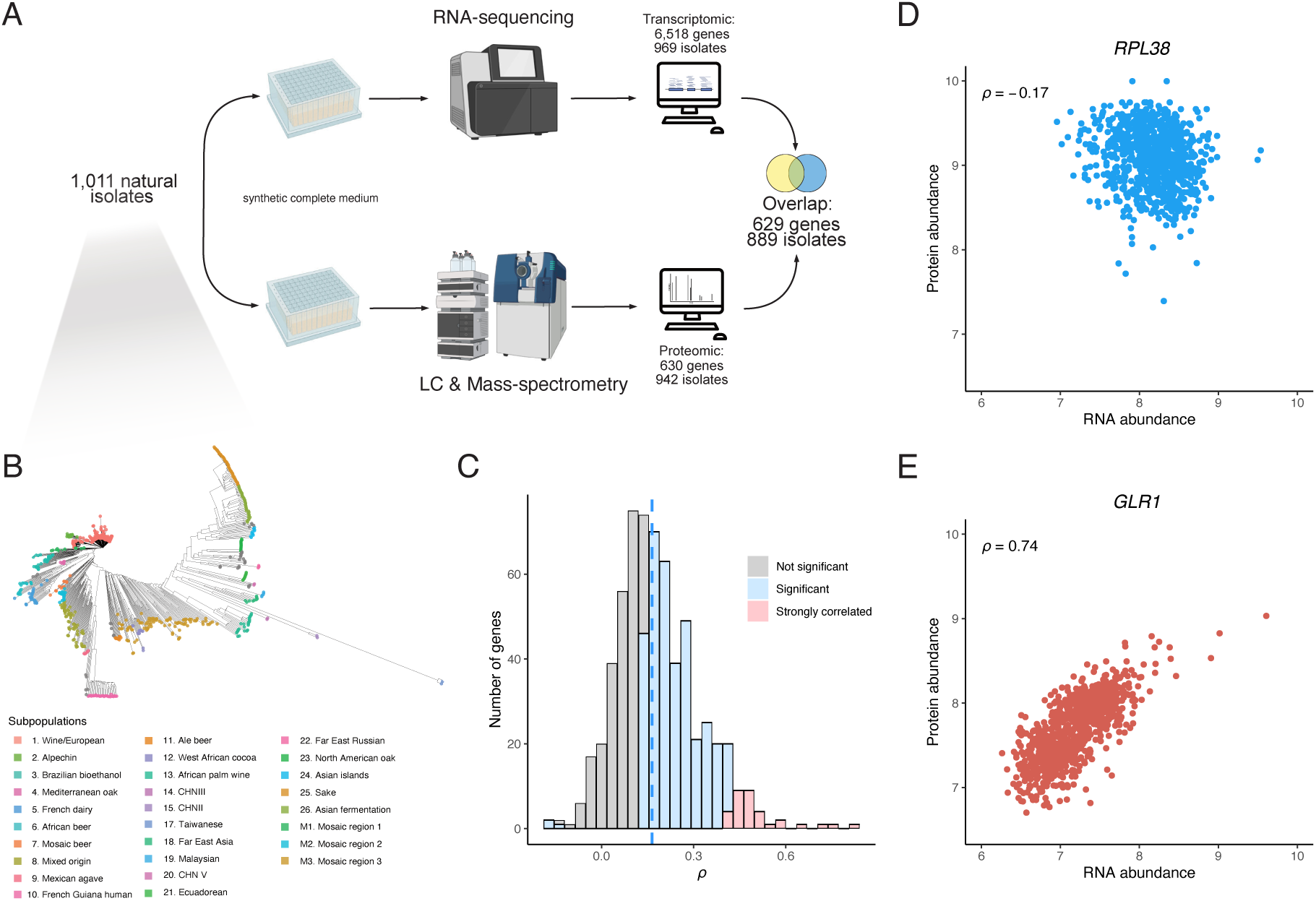
Quantitative proteomes and transcriptomes of a large *S. cerevisiae* population. **A.** The proteomic dataset was generated on isolates grown in synthetic complete (SC) medium with amino acids using a semi-automated sample preparation workflow, and Scanning-SWATH MS (see Methods). The overlap between this dataset and the recently generated transcriptomic dataset on the same population in the same condition^44^ resulted in 629 protein/transcript abundances across 889 isolates. **B.** Phylogenetic trees of the isolates used in this study. Colors correspond to previously defined subpopulations^43^. **C.** Gene-wise correlation coefficients (Spearman correlation test) between the proteome and the transcriptome. **D.** and **E.** mRNA-protein within-gene correlation across isolates for the *RPL38* and *GLR1* genes (*ρ* corresponds to the Spearman correlation coefficient with p-values of 4.8x10^-7^ and 2.3x10^-153^, respectively).

To characterize the quantified proteins in our study, we first compared the level of transcription of both the identified and unidentified proteins. Low abundance transcripts are less likely to be quantified by proteomics as compared to high abundant transcripts (Figure S1D). Indeed, 489 out of 629 consistently quantified proteins fall into the 20% highest transcribed genes (n = 1,304). In total, 537 out of 629 quantified proteins were found in the two highest abundance deciles as defined in a recent yeast protein abundance meta-analysis^49^ (Figure S1E). Overall, proteins related to essential genes and involved in molecular complexes were both significantly enriched in the set of proteins quantified by Scanning SWATH (odd-ratio = 3.5 and 2.2 respectively, Fisher’s exact test, p-values < 2.2x10^-16^)^50–52^. Function-wise, we found that metabolism-related genes were overrepresented among the 629 genes included in our study (Table S4).

We then investigated the level of variation in protein abundance by calculating the coefficient of variation (CV) for each protein using the non-normalized dataset. We found an average CV of 31%, varying between 12% and 98% and one high outlier reaching 300% (PDC5, a pyruvate decarboxylase). The precursor-level CVs across quality control samples (15.15%) were much lower than the precursor-level CVs across the natural isolate samples (34.21%), confirming that a biological signal was observed across the isolates (see Methods). Gene set enrichment analyses (GSEA) were performed using the CVs and significant enrichment of genes related to amino acid metabolism, respiration or pyruvate metabolism was found for proteins with a high CV, indicating that they vary the most (Table S5). By contrast, proteins with a low CV were significantly related to genes involved in tRNA aminoacylation or protein degradation.

### Transcript and protein abundances are weakly correlated at the gene level across isolates

As proteomes and transcriptomes were obtained using the same growth media, our dataset allowed us to characterize the different types of correlation between mRNA and protein abundance across a natural population. We first determined the across-gene correlation, *i.e.* the concordance between protein and transcript abundance for each isolate, and found a very high correlation (median *ρ* = 0.53, interquartile range of 0.06, Figure S2), which is consistent with what was previously described^15–21, 24, 26, 27^. We next computed the correlation between the protein and mRNA normalized abundance for each gene across the 889 natural isolates (Figure 1C-D-E, Table S6). While the across-gene correlation levels were in line with previous explorations, we found an overall very low within-gene correlation level (median *ρ* = 0.165, interquartile range of 0.17). This value is much lower than the one determined with smaller samples in mice (approximately 0.25)^6, 40^ and in human healthy tissues (0.35 and 0.46)^19, 33^, but it is in line with what was found in human lymphoblastoid cell lines (0.14)^15^. For a total of 385 out of the 629 quantified proteins, the level is significantly correlated with RNA level (Bonferroni corrected p-value < 0.05). Out of these 385 proteins, only 3 show a negative correlation: Rps13, Asc1 and Rpl38 (Figure 1D), all ribosomal related proteins. This observation is consistent with previous surveys pointing out that some ribosome-related proteins are negatively correlated with their cognate transcripts^12, 19^. But overall, this correlated set of 385 proteins/transcripts is significantly enriched of genes related to several metabolism pathways (Table S7). Moreover, the most strongly correlated set of proteins/transcripts (n = 33) show functional enrichment of genes related to mitochondrial respiration (Table S8) (see Methods). Interestingly, it points out that this specific pathway has similar gene regulation at both levels. Finally, we observed that four genes with very high mRNA-protein correlation were located outside the main correlation index distribution (Figure 1C). These genes all have correlation coefficients greater than 0.6: *SFA1* (alcohol dehydrogenase), *HBN1* (unknown function), *GLR1* (glutathione oxidoreductase, Figure 1E) and *YLR179C* (unknown function). Such a high correlation clearly points to common regulatory mechanisms and genetic bases underlying the two levels of variation, as we have seen below.

### Gene expression is more constrained at the proteome level

By combining these proteomic and transcriptomic datasets, we are in a position to simultaneously explore and compare the variation of these two gene expression layers at the population level. We therefore computed the absolute *Log2(fold change)* value (*i.e., |Log2(FC)|*) for each gene in each pair of isolates and found that this value is 32% lower on average for the proteome (Figure 2A), suggesting that protein abundance is less variable and more constrained than mRNA abundance. Furthermore, a higher correlation was observed between proteomes (*ρ* = 0.92) compared to transcriptomes (*ρ* = 0.83) (Figure 2B). Finally, the variance observed for each gene was lower for the proteomic data (Figure S3A) and the Euclidean distances between each isolate were smaller when computed with the protein abundance dataset (Figure S3B). Overall, these observations reflect and highlight the presence of a global post-transcriptional buffering of the transcriptome variations.

**Figure 2.**
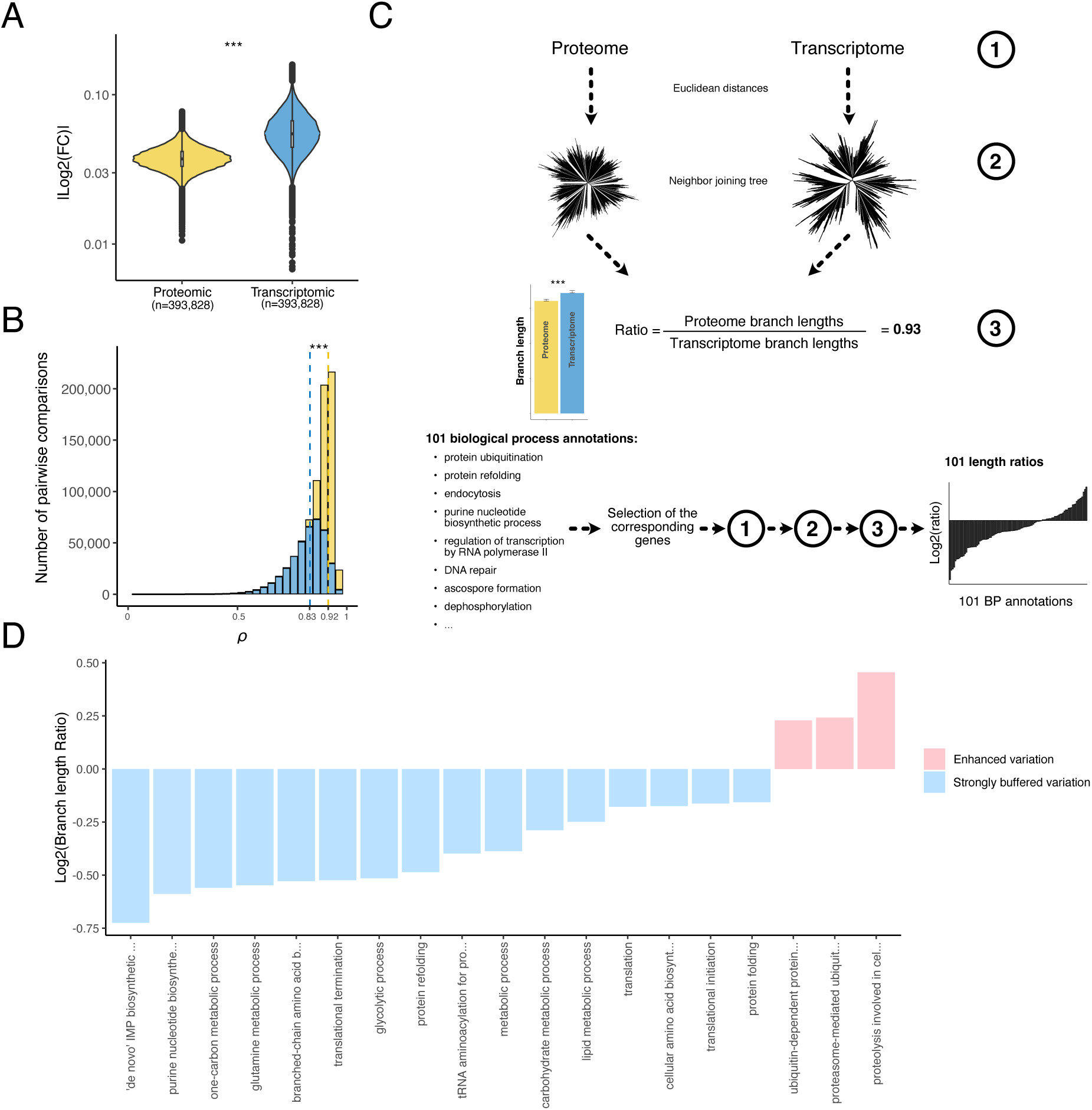
Detection and functional description of the post-transcriptional buffering. **A.** Median | *log2(fold changes)* | computed in each isolate pairwise comparison using both proteomic and transcriptomic data (*** = Wilcoxon test, p-value < 2.2x10^-16^) (see Methods). **B.** Correlation coefficients from the isolate pairwise comparisons using both protein and transcript abundance (*** = Wilcoxon test, p-value < 2.2x10^-16^). The dotted lines correspond to the median correlation index for the proteomic (yellow) and transcriptomic (blue) data. **C.** Cellular functions that are preferentially affected by post-transcriptional buffering. Briefly, using either the proteome and the transcriptome abundances -1- we constructed expression-based neighbor joining trees -2- and compared the total sum of the branch lengths. We computed a ratio -3- defined by the proteome total branch lengths divided by the transcriptome total branch lengths. Using all the genes, this ratio was equal to 0.93 (overall, the expression evolution is more constrained at the proteome level). We performed the same procedure using subsets of genes corresponding to 101 biological process annotations. The biological processes displaying a ratio lower than 0.93 and a significant difference in terms of branch lengths (see Methods) were considered as strongly buffered. The biological processes displaying a ratio higher than 1 and a significant difference in term of branch lengths had an enhanced abundance variation at the proteome level. **D.** Biological processes detected as strongly buffered or with an enhanced variation using the procedure detailed in C.

Despite recurrent observations^45, 53–56^, the post-transcriptional buffering phenomenon remains largely functionally uncharacterized and poorly understood. We sought to better understand this phenomenon at the genetic level by examining the cellular functions that tended to be most affected by post-transcriptional buffering. Briefly, we constructed neighbor-joining trees using the proteome or transcriptome Euclidean distances between each isolate (Figure 2C and see Methods)^56^. Total branch length was used as a measure of expression variation and evolution at the species level. We then calculated the ratio between the lengths of the proteome and transcriptome tree branches to quantify the strength of the post-transcriptional buffering phenomenon. The lengths of branches from the proteome-based tree were shorter than those from the transcriptome-based tree, resulting in a length ratio of 0.93 (Figure 2C, Figure S3C). This observation is consistent with the differences in Euclidean distances observed previously (Figure 2B). We then applied the same procedure to 101 sets of genes, representing central biological processes obtained from a reduced list of gene ontology (GO) annotations (Table S9). We found that a total of 16 sets display a ratio lower than 0.93 and a significant difference between the proteome and transcriptome branch lengths, meaning that these sets are strongly affected by the phenomenon of post-transcriptional buffering (Figure 2D, Table S10). Interestingly, 6 out of the 16 sets include genes with functions related to protein production and maturation (Figure 2D), highlighting that the evolution of the cellular machinery involved in protein production and maturation is highly constrained. The other set of genes are related to several metabolism processes and detected as strongly buffered, despite being highly variable in the proteomic data (Table S5). This observation could be due to the fact that metabolism-related genes are among the genes with the greatest variation in mRNA abundance at the species level^44^. This variation is largely attenuated at the proteome level but remains important, reflecting differences in metabolic preferences within the population. Moreover, we also found 3 sets with a ratio higher than 0.93 and a significant difference between the proteome and transcriptome trees, which means that the expression variation of these genes is greater at the proteome level (Figure 2D, Table S10). Interestingly, all of them are related to protein catabolism, highlighting a difference in post-transcriptional mechanism for this specific functional category. Taken together, these results provide new insights into post-transcriptional buffering as well as its functional impact.

### Architecture of the proteome landscape

Using these datasets, we then sought to understand the main determinants shaping the proteome architecture at the population level. The *S. cerevisiae* yeast species exhibit a clear population structure, which potentially can impact the proteome landscape^43^ (Table S1). We performed a principal component analysis (PCA) with the protein abundance data and found that no clear grouping emerged from the subpopulations when plotting together the 6 first principal components (Figure S4A-B-C). The same results were observed for transcriptomes (Figure S4D-E-F). To confirm this, we also computed the Euclidean distance across transcript and protein levels between every pair of isolates and used these to construct a neighbor-joining tree (Figure 3B-C). We observed that none of the subpopulations present in the genetic-based tree merged in either the proteome- or transcriptome-based tree (Figure 3A-B-C). Together, these results highlight that population structure does not impact transcriptomes and proteomes in the *S. cerevisiae* species.

**Figure 3.**
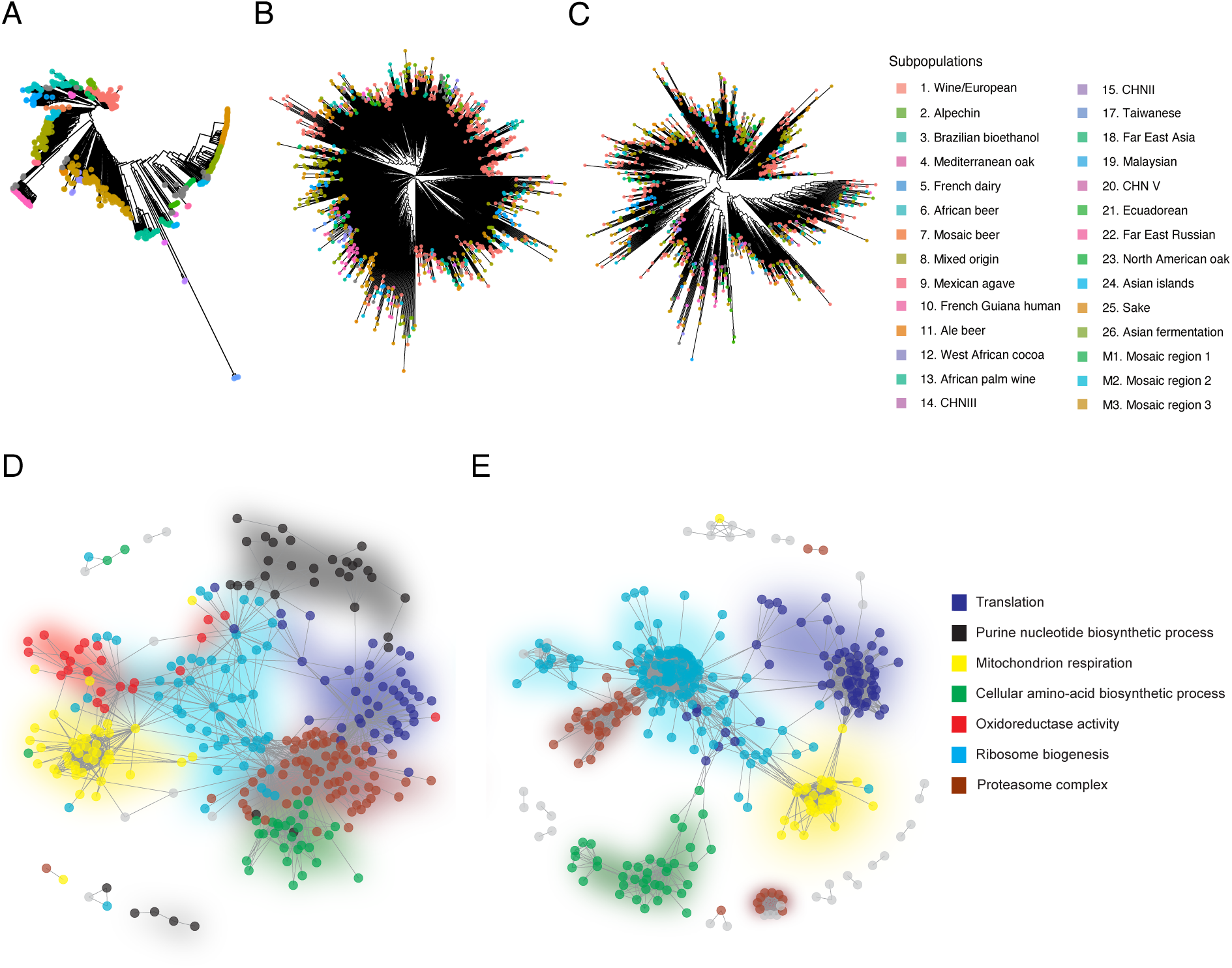
Co-expression network is a major determinant of the proteome organization while the population structure is not. **A-B-C**. Comparison between the phylogenetic tree (A) obtained using the bi-allelic SNP (as in Peter et al. 2018) and the trees obtained from the Euclidean distances based on protein (B) or transcript (C) abundance. Colors correspond to the subpopulations. **D-E**. Cellular co-expression network computed with WGCNA using proteomic (D) or transcriptomic (E) data. Colors represent the cellular pathway detected for each co-expression module.

One potential determinant of the proteome organization could be related to co-expression networks that strongly influence the coordination of gene expression or various cellular processes. Using Weighted Gene Co-Expression Network Analysis (WGCNA)^57^ on the normalized protein abundance data, we detected seven co-expression modules (Figure 3D, Table S11). Each of these modules corresponds to a specific biological function (Table S12, Figure S5) and encompasses between 38 (*Cellular amino acid biosynthetic process*) and 114 (*Ribosome biogenesis*) genes. Interestingly, very similar modules were found applying the same procedure on the mRNA normalized data. Five co-expression modules were detected (Figure 3E, Figure S6, Table S13, Table S14), and all of them were detected in the seven proteomic modules, suggesting that co-expression patterns recapitulate central cell functions are conserved across the two expression layers (Figure 3D-E).

### Insight into subpopulation-specific protein expression

We further wanted to explore and determine the presence of subpopulation-specific signatures. We therefore sought to identify differential protein expression patterns by comparing each clade to the rest of the population and we detected a total number of 1,129 differentially expressed proteins (DEPs) (corresponding to 465 unique proteins, Figure S7, Table S15). An average of 59 DEPs was found per clade, ranging from 218 for the Wine clade to 0 for wild Asian clades represented by a small sample (*e.g.* CHN, Taiwanese and Far East Russian) (Figure S8A). Several DEPs were adequately related to the ecological origin of the different subpopulations. For example, several subpopulations related to alcoholic fermentation show overexpression of alcohol dehydrogenases, such as ADH4 in Wine and Brazilian bioethanol clades as well as ADH3 in the Sake subpopulation. In the French Dairy subpopulation, we also observed an underexpression of SEC23, a GTPase-activating protein involved in the COPII related vesicle formation, which could reflect an adaptation to this secretory pathway to the cheese-making environment^58^. Overall, these observations suggest that domestication and more generally, ecological constraints are drivers of the proteomic landscape evolution in a natural population. We then performed GSEA based on differential expressed proteins in each subpopulation and found significant enrichments for various biological processes (Figure 4A, Table S16). Many enriched functional categories were associated with respiration related genes (*e.g.* “*respiratory electron chain transport*”). Interestingly, we observed that while most wild clades (8 out of 13) tend to have overexpression of respiration-related proteins, these are underexpressed in domesticated subpopulations (5 out of 7).

**Figure 4.**
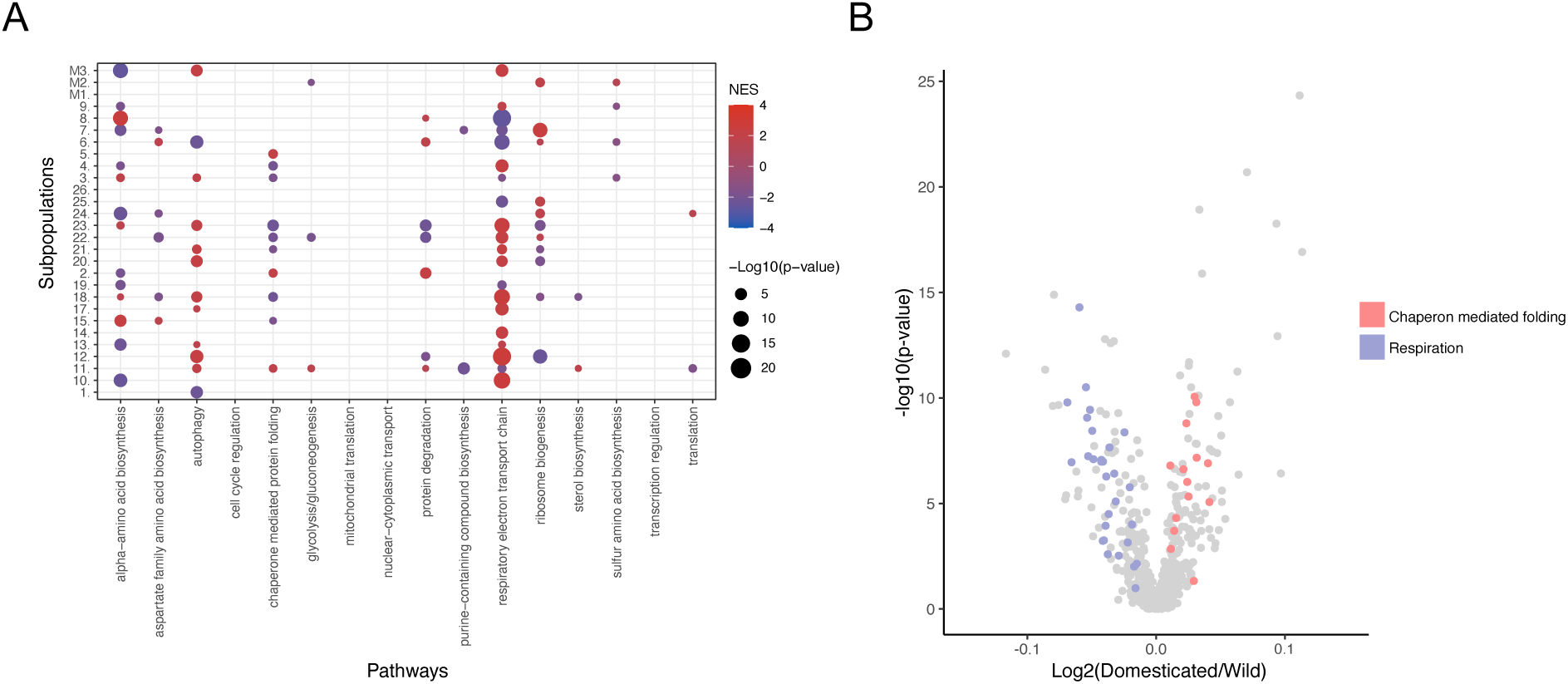
DEPs reveal domestication- and subpopulation-specific metabolic adaptation. **A.** GSEA results on the DEPs (using 16 broad functional annotations from *44*) of each subpopulation. Colors represent the normalized enrichment score (NES): Red – overexpression, blue – underexpression in subpopulation. **B.** Volcano plot of the comparison between wild and domesticated isolates. Colors highlight the genes belonging to two functional annotations related to chaperon mediated folding and respiration.

We therefore further explored the impact of domestication on the proteome at the population level. Using the same DEP detection method, we assessed the proteome differences between the domesticated and wild isolates^43^ and found a total of 133 DEPs (Table S17). Among these proteins, other alcohol dehydrogenases such as SFA1 and ADH3 were highly abundant in domesticated isolates. A GSEA performed on this set of DEPs clearly shows an enrichment of underexpressed respiration-related proteins in domesticated clades (Figure 4B, Table S18). Unlike wild isolates, domesticated isolates were selected for fermentation purposes, likely leading to this specific signature. This observation is in line with the previous finding pointing out that the switch from a preference between respiration and fermentation is one of the hallmarks of domestication in yeast^59^. In addition, significant enrichment of the functional category “*chaperon mediated protein folding*” points to overexpression of this set of proteins in the domesticated isolates (Figure 4B), which may be an adaptative response to long-term exposure to ethanol, known to induce protein denaturation^60^. By performing the same analysis on transcriptomic data (Figure S8B, Table S17), similar results, showing overexpression of respiration-related genes in domesticated clades, were obtained (Table S19).

### The genetic bases of protein abundance at the population scale

To uncover the genetic origins of the proteome variation at the population-scale, we performed genome-wide association studies (GWAS) and considered both SNPs and CNVs that were characterized previously^43^. We focused on isolates for which both proteomic and transcriptomic data were available, resulting in a set of 889 isolates. In this population, a total of 84,633 SNPs and 1,019 CNVs were considered, with a minor allele frequency higher than 5%. We performed GWAS using the raw protein abundances of the genes for which we have both levels of expression (*i.e.*, 629 genes). Overall, we detected a total of 598 SNP-pQTL after colliding SNP affected by linkage disequilibrium (R2 > 0.6), and 4,528 CNV-pQTL corresponding to 501 and 520 loci and affecting 300 and 93 genes, respectively (Figure 5A-B, Table S20, Table S21, data file 1).

**Figure 5.**
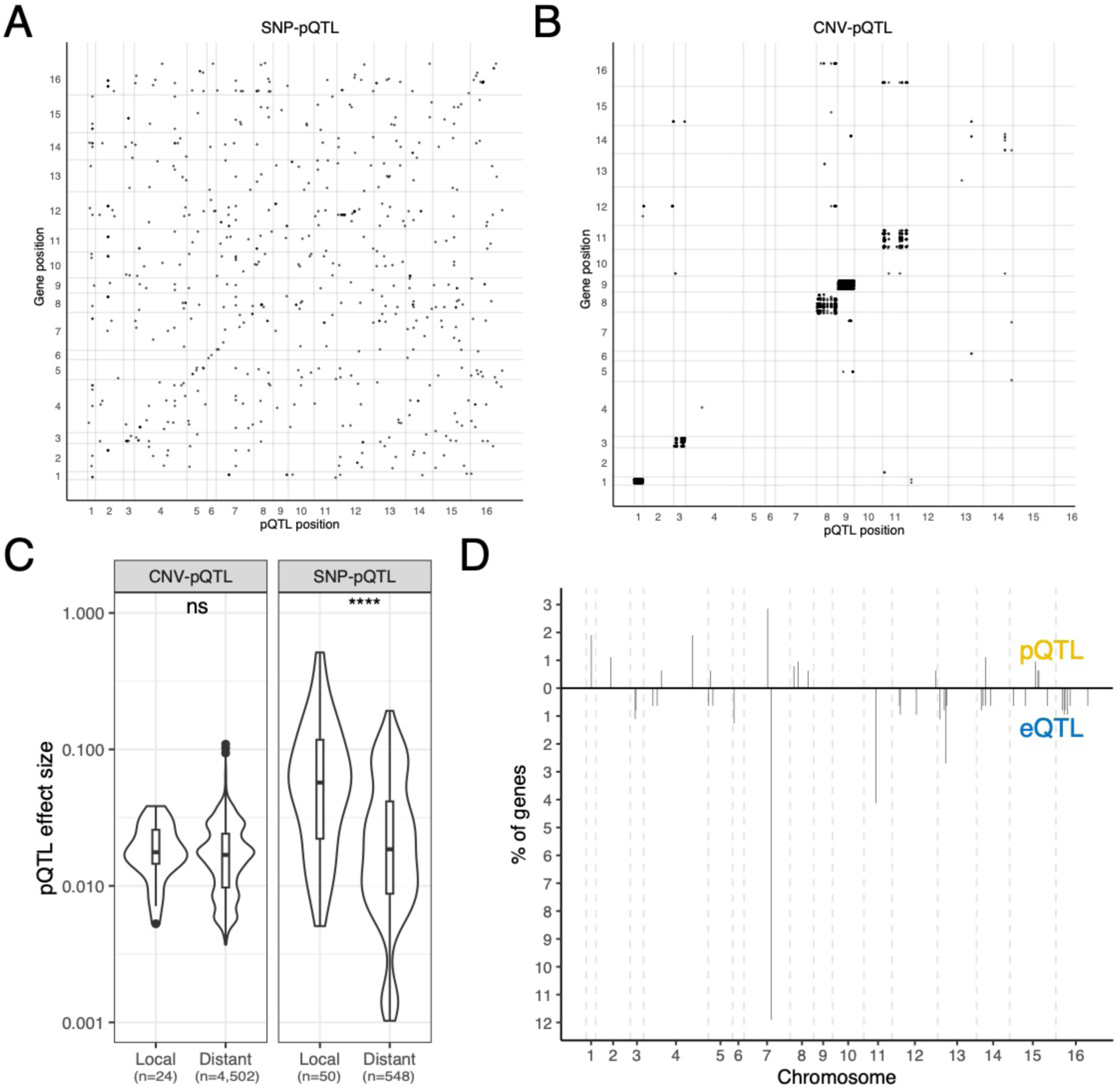
SNP- and CNV-pQTL detection highlights strong differences in the genetic origin of transcript and protein abundance. **A-B**. Map of the SNP- (A) and CNV- (B) pQTL. The x-axis is the QTL position on the genome and the y-axis the position of the affected gene on the genome. The x and y-axis numbers represent the 16 chromosomes of *S. cerevisiae*. **C.** Effect size difference between the local and distant pQTL for the SNP (p-value= 6.2x10^-8^) and CNV pQTL (p-value=0.38). **D.** Distribution of the SNP-pQTL and SNP-eQTL hotspots along the genome. The y-axis represents the percentage of the 629 genes that by each hotspot (defined as a 20 kb window containing 4 or more distinct SNP).

Among the SNP-pQTL, 8% (n = 50) were local-pQTL, showing that regulation of protein abundance is primarily achieved through *trans* regulation. This fraction is consistent with previous exploration in yeast^61^ and lower than what is usually found at the transcriptome level^44, 62^. Nonetheless, we observed that the local SNP-pQTL have a higher effect size compared to *trans* SNP-pQTL (Figure 5C) and tend to be located near the transcription starting site of the gene (Figure S9). We found no strong SNP-pQTL hotspots, suggesting that most of the distant pQTL are evenly distributed throughout the genome (Figure 5D).

In contrast, CNVs impacting protein abundance had a biased location on the chromosomes 1, 3, 8, 9 and 11 (Figure 5B). Out of 4,528 CNV-pQTL, a total of 4,303 were located on these chromosomes and affected a gene on their respective chromosome. This observed bias is due to the presence of aneuploidies on these chromosomes in our population^43^. These CNV-pQTL have also a higher impact on the protein abundance variation compared to the other CNV-pQTL, suggesting that aneuploidies represent a major source of proteome variation at the population level (Figure S10). Only 24 local CNV-pQTL out of 4,528 were detected, and no significant effect size between local and distant CNV-pQTL was found (Figure 5C).

We then looked at the extent to which the genetic bases of protein abundance are common with those underlying the abundance of transcripts. We performed GWAS using the transcriptomic dataset and detected 596 SNP-eQTL and 4,877 CNV-eQTL (Figure S11, data file 2), which is of the same order of magnitude as the GWAS proteome results. Surprisingly, the overlap between the SNP-pQTL and the SNP-eQTL is very low, with only 3.6% of shared SNP-QTL (n = 22). Interestingly, 18 out of 22 were related to local regulation, meaning that 36% of the local SNP-pQTL (18 out of 50) also impact the cognate transcripts of their target protein. This observation is consistent with previous findings showing that the common regulation between mRNA and protein abundances is mainly related to local regulation^6, 40^. Overall, we observed that genes with a strong correlation between transcript and protein abundance, such as the top four most correlated genes previously mentioned (*SFA1*, *HBN1*, *GLR1* and *YLR179C*), tend to have a shared pQTL and eQTL (Figure S12). Additionally, we found that the SNP-pQTL distribution across the genome did not match the SNP-eQTL distribution, where a QTL hotspot could be detected around the *CTT1* gene^44, 63^. The reasons for the weak overlap are likely multifactorial, but protein-specific regulation, such as protein degradation, may play a central role. We sought to confirm this by looking at the average protein turnover^45^ of the proteins with and without overlapping pQTL and eQTL (Figure S13A, see Methods). We found that proteins, for which an overlap between pQTL and eQTL was detected, show a lower turnover rate compared to the other proteins. Consistently, the half-life of proteins with an overlapping SNP-QTL was higher than the rest of the proteome (Figure S13B). This observation suggests that protein degradation is probably involved in the large differences observed between the genetic origins of mRNA and protein abundance.

In contrast, the overlap between the two sets of CNV-QTL is much higher, as 3,097 QTLs were shared between the transcriptome and proteome, *i.e.*, approximately 68% of the CNV-pQTL. However, these shared CNV-QTLs are all aneuploidy-related CNVs, suggesting that the effect of aneuploidies is persistent through the expression layers^45^. None of the non-aneuploidy CNV-QTL (22 CNV-eQTL and 216 CNV-pQTL) were shared. Together, our results highlight that the genetic bases underlying population-level protein abundance are very distinct from those underlying mRNA abundance.

## Discussion

Quantifying transcripts and proteins expressed in a large natural population is fundamental for having a better understanding of the genotype-phenotype relationship. In this study, we have quantitatively analyzed the proteome of 942 natural isolates of *S. cerevisiae*, allowing in-depth exploration of protein abundance and precise characterization of the genetic origins of its variation at the species level.

The *S. cerevisiae* species is characterized by a complex population structure, with domesticated and wild subpopulations^43^. Structured populations are also observed in a large number of other species, such as humans, and their impact on the proteome remains unexplored. In our dataset, the population structure had no significant impact on the proteomic landscape. This observation is consistent with previous results obtained with the transcriptomes of *S. cerevisiae* isolates^44, 64^. In fact, most subpopulations are characterized by specific signatures related to a small set of genes but not to a general pattern.

This dataset allowed us to have better insight into the architecture of the species-wide proteome variation. First, we found that the co-expression network captures main biological functions and is globally conserved across the species. Second, we detected differential protein expression signatures specific to subpopulations, reflecting an adaptation to specific ecological conditions, such as domesticated environments. Similar expression signatures can be also observed using transcriptomic data^44, 65^, highlighting that gene expression plasticity at both levels is a key mechanism of environmental adaptation.

The species-wide proteomes and transcriptomes obtained in the same condition represent a unique opportunity to compare the gene regulation at both levels. The overall agreement between protein and transcript within each isolate appears to be high and this in the whole population, showing again that very abundant transcripts generally lead to very abundant proteins and vice versa^15–21, 24, 26, 27^. However, our data allow for the first time to have an accurate estimation of the correlation per gene at the population level and we found that this gene-wise correlation is very weak with a median of 0.165, which is lower than most previous estimates based on much smaller human and mice populations^6, 19, 33, 37, 40^. Consistent with this result, genome-wide association studies also highlighted that SNPs related to variation in protein (pQTL) and transcript (eQTL) levels poorly overlap (3.6%), with mostly common local QTL. This result is consistent with one of the first eQTL/pQTL comparisons^61^ but unlike other studies, showing a higher overlap^66^. However, we should emphasize that we were not able to map the genetic basis of the entire *S. cerevisiae* proteome and therefore the eQTL/pQTL overlap might be biased and underestimated.

Mechanistically, our results suggest that the regulation of protein degradation has an impact on the variation of the proteome, and therefore on its genetic basis. Proteins with a high turnover rate will be more affected by proteome-specific regulation and will therefore show a weaker correspondence with the transcriptome. Conversely, proteins with a low turnover rate are more likely to be impacted by variation in transcript abundance. They will therefore likely reflect variation in mRNA abundance.

Although mass spectrometers are highly sensitive, it should be noted the limitation that proteomic methods are biased towards quantification of highly abundant proteins. Indeed, the fraction of the proteome quantified constitutes the vast majority of the total proteomic mass of a cell and is enriched for essential genes as well as in genes most connected in functional networks. Our dataset captures many of the fundamental processes. Yet, results related to low abundant proteins are missed by this approach.

Overall, our study clearly highlights that the dependency between transcript and protein levels is complex, pointing to the importance of post-transcriptional regulation of protein abundance. Proteome and transcriptome are indeed two distinct layers of gene regulation, which need to be further explored to understand the genotype-phenotype relationship. As gene function is ultimately executed by the proteome, while mRNA is the messenger, more proteomic approaches will be needed to create a better understanding of the phenotypic diversity. Our study provides a first species-wide insight into the genetics that underlies both proteome and transcriptome diversity in a natural population.

## Methods

### Cultivation of library for proteomics

The yeast isolate collection was grown on agar containing synthetic complete medium (SC; 6.7 g/L yeast nitrogen base (MP Biomedicals, Cat#114027512-CF), 20 g/L glucose, 2 g/L synthetic complete amino acid mixture (MP Biomedicals, Cat#114400022)). After 48 h, colonies were inoculated in 200 µL SC liquid medium using a Singer Rotor and incubated at 30 °C overnight without shaking. These pre-cultures were then mixed by pipetting up and down, and diluted 20x by transferring 80 µL per culture to deep-well plates pre-filled with 1.55 mL SC liquid medium and one borosilicate glass bead per well. Plates were sealed with a permeable membrane and grown for 8 h at 1000 rpm, 30°C to exponential phase. The optical density at harvest was measured using an Infinite M Nano (Tecan). Per culture, 1.4 mL of cell suspension were harvested by transferring into a new deep-well plate and subsequent centrifugation (3,220 x g, 5 min, 4°C). The supernatant was removed by inverting the plates. Cell pellets were immediately cooled on dry ice and stored at −80°C.

### Sample preparation

Samples for proteomics were prepared as previously described^42, 45, 47^. In brief, samples were processed in 96-well format, with lysis being achieved by beat beating using a Spex Geno/Grinder and 200 µL of lysis buffer (100 mM ammonium bicarbonate, 7 M urea). Samples were reduced and alkylated using DTT (20 µL, 55 mM) and iodoacetamide (20 μL, 120 mM), respectively, diluted with 1 mL 100 mM ammonium bicarbonate, and 500 µL per sample were digested using 2 µg Trypsin/LysC (Promega, Cat#V5072). After 17 h of incubation at 37°C, 25 µL 20% formic acid were added to the samples, and peptides were purified using solid-phase extraction as described previously^47^. Eluted samples were vacuum-dried and subsequently dissolved in 70 µL 0.1% formic acid. An equivoluminal pool of all samples was generated to be used as technical controls (QCs) during MS measurements. The peptide concentration of this pool was determined using a fluorimetric peptide assay kit (Thermo Scientific, Cat#23290). Peptide concentrations per sample were estimated by multiplying the optical density recorded at harvest with the ratio between pool peptide concentration and the median at-harvest optical density.

### LC–MS/MS measurements

In brief, peptides were separated on a 3-min high-flow chromatographic gradient and recorded by mass spectrometry using Scanning SWATH^46^ using an online coupled 1290 Infinity II LC system (Agilent) -6600+ TripleTOF platform (Sciex). 5 µg of sample were injected onto a reverse phase HPLC column (Luna^®^Omega 1.6µm C18 100A, 30 × 2.1 mm, Phenomenex) and resolved by gradient elution at a flow rate of 800 µL/min and column temperature of 30 °C. All solvents were of LC-MS grade. The gradient program used 0.1% formic acid in water (Solvent A) and 0.1% formic acid in acetonitrile (Solvent B) and was as follows: 1% to 40% B in 3 min, increase to 80% B at 1.2 mL over 0.5 min, which was maintained for 0.2 min and followed by equilibration with starting conditions for 1 min. For mass spectrometry analysis, the scanning swath precursor isolation window was 10 m/z; the bin size was set to 1/5th of the window size, the cycle time was 0.7 s, the precursor range 400 m/z to 900 m/z, the fragment range 100 m/z to 1500 m/z as previously described^46^. An IonDrive TurboV source was used with ion source gas 1 (nebulizer gas), ion source gas 2 (heater gas), and curtain gas set to 50 psi, 40 psi and 25 psi respectively. The source temperature was set to 450 °C and the ion spray voltage to 5500 V.

### Data processing

The mass spectrometry files were processed following the approach previously described^45^. Briefly, an experimental spectral library obtained using the S288c was filtered to reduce the search space to peptides well shared across the strains. This library was then used with the software DIA-NN^48^ (Version 1.8) and the following parameters: missed cleavages: 0, Mass accuracy: 20, Mass accuracy MS1: 12, scan windows: 6. The option ‘MBR’ was used to process the data. As the peptides selected were not necessarily present ubiquitously in all the strains, an additional step was required to remove false positives (entries where a peptide is detected in a strain where it should be absent). This represents only ∼1% of the total entries of the report.

Samples and entries with insufficient MS2 signal quality (< 1/3 of median MS2 signal) and with entries with Q.Value (> 0.01), PG.Q.Value (> 0.01), Global.Protein.Q.Value (> 0.01), Global.PG.Q.Value (> 0.01) were removed. A similar threshold was applied to Lib.PG.Q.Value and Lib.Q.Value to account for the MBR option used. Non-proteotypic precursors were also excluded. Outlier samples were detected based on the total ion chromatograms (TIC) and number of identified precursors per sample (Z-Score > 2.5) and were excluded from further analysis. Precursors were filtered according to their detection rate in the samples, with a threshold set at 80% of detection rate across all the strains, while precursors with a coefficient of variation (CV) above 0.3 in the QC samples were excluded. The CVs of QCs and wild isolates samples were calculated and had a median CV of 15.15% and 34.21%, respectively (Figure S14; table S22). Batch correction was carried out at the precursor level using median batch correction, which consists in bringing the median value of the precursors in the different batches to the same level. Proteins were then quantified from the peptide abundance using the maxLFQ^67^ function implemented in the DIA-NN R package. The resulting dataset consists of 630 proteins for 942 strains. We imputed the missing value for further exploration using the KNN imputation method from the *impute* R package^68^.

### Combination of transcriptomic and proteomic data

Unless specified, all the analysis performed below were conducted using R version 4.1.2. The transcriptomic data was generated previously^44^. We used the log2 transcript per million (TPM) data, where the overlap with proteomic data was encompassing 629 genes across 889 isolates, for the genome wide association studies (see later for the method). For the exploration of gene expression variation, subpopulation related DEG and gene expression network, we used the variance stabilized data obtained directly from the log2 TPM data. In this case one gene was removed from the analysis and the reference strain data was not considered, which resulted in an overlap of 628 genes across 888 isolates. To only focus on real expression variation difference between the expression layers, we normalized the proteomic and transcript abundance using quantile normalization. Unless specified, all the analyses described below use the quantile normalized transcriptomic and proteomic data. We recomputed the raw protein abundance coefficient of variation (CV) of each gene by dividing the standard deviation by the mean (using the non-normalized abundance) and transformed it to a percentage. Based on the CV, we performed a functional exploration by gene set enrichment analysis (GSEA)^69^ using the *fgsea* R package^70^ for the gene ontology annotation^71, 72^ to detect cellular pathways with a conserved regulation across the population. The within- and across-gene mRNA-protein correlation was performed for each gene or each isolate using a Spearman correlation test. We selected the genes with a mRNA-protein correlation index higher than 0.42 (> 95% percentile) and performed gene ontology (GO) enrichment analysis using the biological process (BP) database using the *topGO* R package^73^. For the GO analysis looking at the functional enrichment present in the 630, the gene list reference was the genes encompassed in the transcriptomic data^44^. The others GO analyses used the 628 genes as the reference list. All the others GO analyses were performed using the same procedure, unless specified.

### Expression variation exploration

We measured the strength of protein and transcript abundance variation using several methods. We computed an absolute transformed *Log2(fold change)* value (*|Log2(FC)|*) where in each isolate pairwise comparison (ex: strain A vs strain B) and for each gene, we performed:

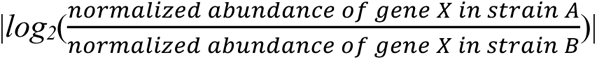

Briefly, the more this value increases, the more different is protein abundance between two isolates for a specific gene. We also computed a pairwise spearman correlation between the isolates using the normalized proteomic and transcriptomic data. We also gathered the Euclidean distances between the expression profiles of each isolate, as well as the gene expression variance per gene.

We explored the post-transcriptional buffering phenomenon using an approach based on the computation of expression trees^56^. First, on both protein and transcript normalized abundances, we constructed a neighbor-joining tree based on the Euclidean distance between each isolate. We computed the total branch length of these two trees and created a ratio of the proteome tree length on the transcriptome tree length. The ratio was equal to 0.93 which is line with the difference in Euclidean distance between the transcriptome and proteome. We performed 100 bootstrapping tests and used the resulting branch lengths to test the difference between the proteome and the transcriptome tree. We sought to check if some cellular pathways tended to be more affected by the post transcriptional buffering phenomenon. To do so, we gathered a reduced biological process GO annotation by computing the similarity between each GO term using the *rrvgo* R package and the ‘Resnik” method^74^. We discarded terms that are at least 50% overlapping with another term and the terms encompassing no more than 5 genes, which resulted in a list of 101 terms. For each of these terms, we performed the same tree exploration, but this time with the genes encompassed by each term. We obtained therefore 101 tree length ratios. We selected the terms displaying a ratio lower than 0.93 or higher than 1, and for which the total branch length between the proteome and the transcriptome was significantly different after 10 bootstrapping steps (Bonferroni corrected Wilcoxon test p-value < 0.001).

### Transcriptome and proteome landscape exploration

We sought to check if the genetic structure of the population had an impact on the transcriptome and proteome structure. We obtained the genetic distances from^43^ between pairs of isolates and compared them to the pairwise isolate correlation (Spearman correlation test) obtained with the normalized transcript or protein abundances. We also used both normalized protein and mRNA abundance data to perform principal component analysis (PCA) using the *prcomp* function from the *stats* R package. For the 2 PCA (transcriptomic and proteomic), we plotted the 6 first principal components (PC) together (PC1-PC2, PC3-PC4 and PC5-PC6) and looked for eventual grouping according to the subpopulation as defined previously^43^. We then computed a Weighted Gene Co-Expression Network Analysis (WGCNA) using the *WGCNA* R package^75^ to detect co-expression module in both mRNA and peptide normalized abundance. To do so, we generated a Topological Overlap Matrix (TOM) using the *blockwiseModules* function. The TOM were calculated based on a signed adjacency matrix with the power of 9 for the mRNA abundance data and 5 for the peptide abundance data. The *blockwiseModules* automatically detected the co expression modules by generating a clustering from a dissimilarity matrix (1-TOM) using the following option: *detectCutHeight* = 0.995; *minModuleSize* = 30. This resulted in the detection of 5 and 7 transcriptome and proteome modules respectively. We computed an overrepresentation analysis for each co-expression module with the GO terms as annotation and using the *mod_ora* function from the *CEMiTool* R package^76^ and used the most representative GO terms as the final annotation for each detected module. The two co-expression networks were generated for plotting by computing an adjacency matrix from the TOM matrix (generated previously) and ultimately plotted using the *ggnet2* function from the *GGally* R package.

### Transcriptome and proteome differentially expressed gene detection

We used the normalized protein abundance to detect subpopulation-specific^43^ differentially expressed proteins (DEPs). The goal was to detect either over- or underexpressed genes by comparing the normalized expression of all the isolates from a subpopulation against the rest of the population using a Wilcoxon test for each gene. The p-value of the test was corrected using a Bonferroni correction with the *p.adjust* function in R. A gene was considered as differentially expressed if the corrected p-value of the Wilcoxon test was below 0.05. We computed as well a log2 transformed fold change (log2(FC)) value for each gene in each subpopulation using the mean expression of the subpopulation divided by the mean expression of the rest of the population. To further characterize the detected DEPs, we performed a functional exploration using GSEA (with the *fgsea* function from the *fgsea* R package) using the log2(FC) value from the DEP exploration as score rankings. In order to have a global view of the pathways that were significantly differentially expressed in each subpopulation, we used the 16 co-expression modules detected and defined previously using the population transcriptome data^44^ as biological function annotations for the GSEA. We performed the same procedure but this time comparing the domesticated against the wild isolate (using the clade wise annotation from^43^). This time, the test was performed on both normalized protein and transcript abundances.

### Proteome and transcriptome genome-wide association studies

We computed GWAS with a linear mixed model-based method as described previously^43, 44^ using FaST-LMM^77^. We performed the GWAS using either the transcriptome log2 transformed TPM data or the protein abundance. For each dataset, we performed two separated GWAS, one based on SNP as genotype, and one based on the CNV as genotype. The SNP GWAS was run with total of 84,633 SNP displaying a minor allele frequency (MAF) > 5% and that were not located in the telomeric regions (< 20kb away from the chromosome ends). The CNV GWAS was run on a total of 1,019 CNV (MAF > 5%). We used the SNP matrix for both SNP and CNV GWAS, thus evaluating the kinship between the isolate to account for the population structure. We set a phenotype-specific p-value threshold using 100 permutation tests where the phenotypes were randomly permutated between the isolates. We use the 5% lowest p-value quantile from these permutation tests to define the significance threshold. We finally scaled the significance thresholds of the CNV GWAS to account for the size difference between the SNP and CNV matrices.

Regarding the SNP GWAS, the detected QTL were filtered to avoid false positives detection due to linkage disequilibrium among the SNP as described previously^44^. This resulted in the filtration of 81 eQTL and 131 pQTL (out of respectively 677 and 729 QTL). The QTL were considered as “local” QTL when they were located 25 kb around their affected phenotype. We also sought to detect QTL hotspots in both transcriptome and proteome GWAS. We defined a hotspot as a concentration of at least 4 QTL in a 20 kb window.

We compared the protein turnover rate^45^ of obtained on 619 proteins encompassed in our dataset to see whether turnover rate had an impact on the overlap between SNP-eQTL and SNP-pQTL. This data comprises protein degradation rates for 1,836 gene across 55 natural isolates. We computed an average turnover rate per gene and used this value to compare the level of protein degradation of the protein with or without an overlapping QTL.

## Supporting information

All_sup_figures

## Acknowledgements

This work was supported by a National Institutes of Health (NIH) grant R01 (GM147040-01) and a European Research Council (ERC) Consolidator grant (772505) to J.S. And E.T. was supported by the PhD Joint Programme CNRS & Weizmann Institute and a fellowship from the medical association la Fondation pour la Recherche Médicale (FDT202204014796). J.S. is a Fellow of the University of Strasbourg Institute for Advanced Study (USIAS) and a member of the Institut Universitaire de France.

