## Supplementary material for "Species-wide quantitative transcriptomes and proteomes reveal distinct genetic control of gene expression variation in yeast": All_sup_figures

**This document includes:**

Supplementary figures S1-11

Supplementary figure legends

**Figure S1**. **Description of the characteristics and the normalization of the population proteome.**

(A) number of isolates encompassed in the proteomic datasets, in the transcriptomic dataset (Caudal et al., submitted) and in the overall population (Peter et al., 2018). The x-axis corresponds to the clades (or subpopulations) as defined previously (Peter et al., 2018). (B, C) Expression values of 5 randomly selected isolates for protein and transcript abundance before (B) and after (C) quantile normalization. (D) mRNA levels of the gene encompassed or not in the proteomic data (*** = p-value < 2.2x10^-16^, Wilcoxon test). (E) Protein levels (as defined in Ho et al., 2018) of the genes encompassed by our proteomic data.

**Figure S2. Across-gene correlation.**

mRNA-protein correlation in each isolate (across-gene correlation). The blue line represents the median (0.53).

**Figure S3. Detection of the post-transcriptional buffering.**

(A) Comparison between the gene-wise protein and transcript normalized abundance variance (*** = p-value < 2.2x10^-16^, Wilcoxon test). (B) Euclidean distances between each isolate using the protein or transcript normalized abundance (*** = p-value < 2.2x10^-16^, Wilcoxon test). (C) Branch length difference between the proteome and the transcriptome-based tree. The error bars correspond to 100 bootstrapping steps. We used the bootstrap values to test if the difference in branch length is significant between the two trees (*** = p-value < 2.2x10^-16^, Wilcoxon test).

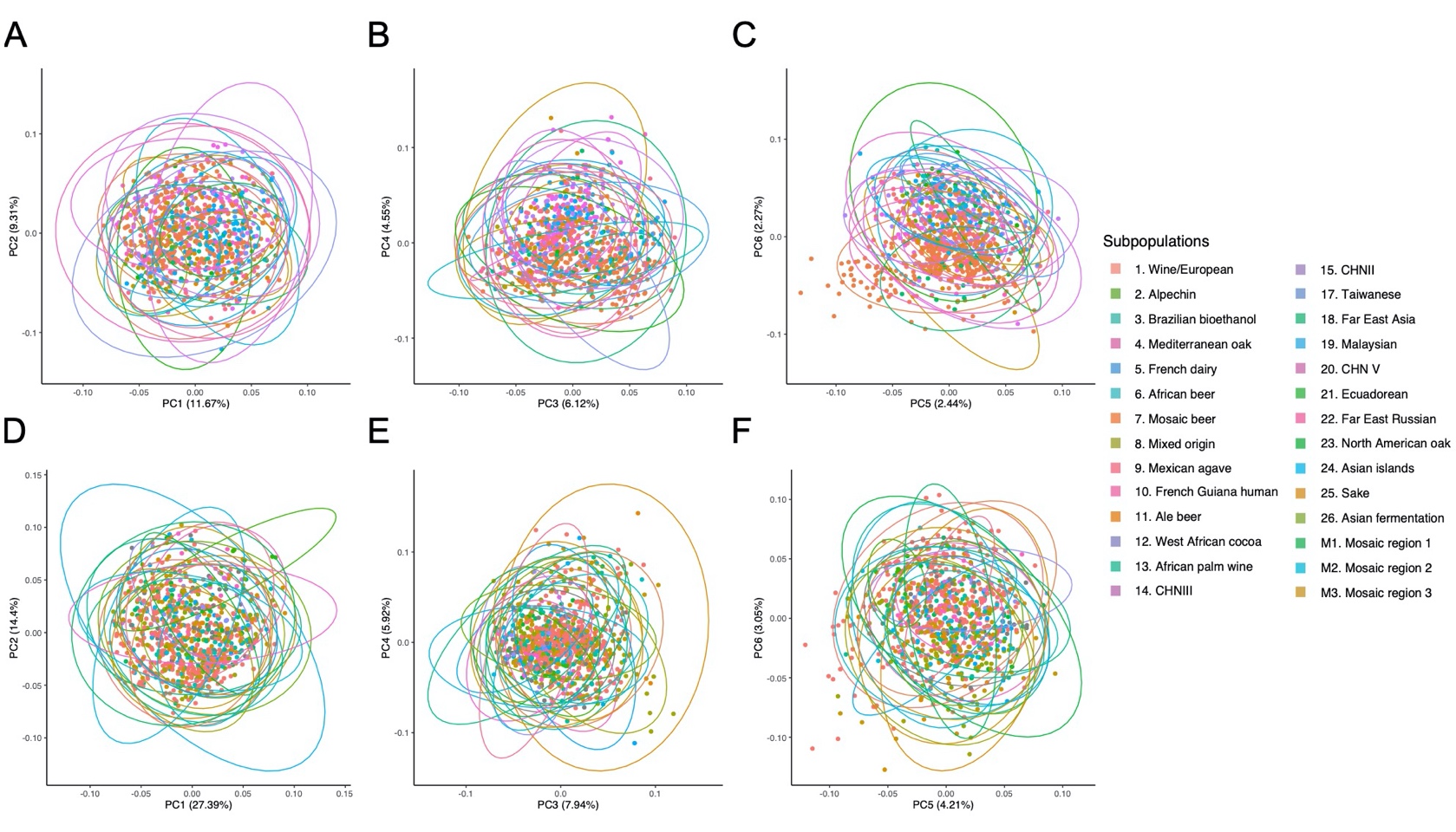

**Figure S4. Population structure is not reflected on the proteome based PCA.**

PCA using protein (A, B, C) or transcript (D, E, F) abundance. The 6 first PC are plotted together, and the colors correspond to the subpopulations (clades).

**Figure S5. Functional exploration of the proteome WGCNA modules.**

Functional enrichment of each co-expression module detected using WGCNA on protein abundance data. The enrichment was performed using the CEMiTool package. The dotted lines on each graph represent the significance threshold.

**Figure S6**. **Functional exploration of the transcriptome WGCNA modules.**

Functional enrichment of each co-expression module detected using WGCNA on transcript abundance data. The enrichment was performed using the CEMiTool package. The dotted lines on each graph represent the significance threshold.

**Figure S7. DEPs detected in each subpopulation.**

Volcano plots for each subpopulation highlighting the DEPs. The blue points correspond to under-expressed gene in a subpopulation while the red points correspond to over-expressed genes.

**Figure S8.** **Number of DEP and differentially expressed transcripts.**

(A, B) Number of proteome (A) and transcriptome (B) DEPs (or differentially expressed transcripts for the transcriptome) in each subpopulation together with the number of isolates in each subpopulation.

**Figure S9. Location of the local SNP-pQTL.**

Distribution of the local SNP-pQTL around the start codon of their target gene. Downstream pQTL correspond to QTL located between the stop codon and 200 bp after the stop codon, upstream correspond to pQTL located between the start codon and 1,000 bp before the start codon.

**Figure S10. Aneuploidy related CNV-pQTL have a higher effect-size than the other CNV-pQTL.**

Difference in effect size between the aneuploidy related CNV-pQTL and the other CNV-pQTL (*** = p-value < 2.2x10^-16^, Wilcoxon test).

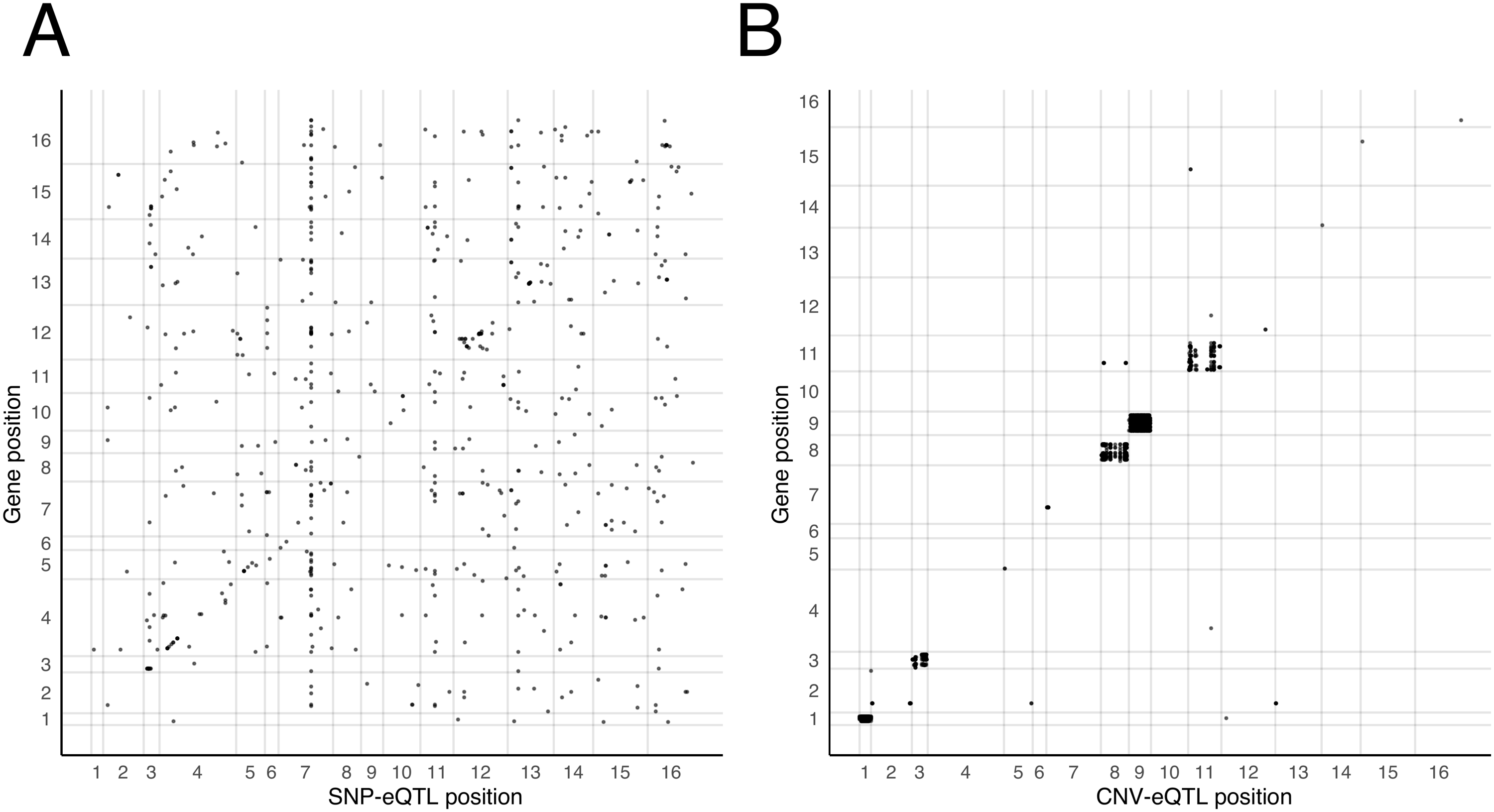

**Figure S11. Genomic location of the SNP- and CNV-eQTL.**

(A, B) Map of the SNP (A) and CNV (B) eQTL. The x-axis is the QTL positions on the genome and the y-axis the position of the affected genes on the genome. The x and y-axis numbers represent the 16 chromosomes of *S. cerevisiae*.

**Figure S12. The genes with an overlapping SNP-QTL tend to have a high within-gene mRNA-protein correlation.**

Within-gene correlation coefficients (Spearman correlation test) between the proteome and the transcriptome. The genes with an overlapping SNP-pQTL and SNP-eQTL are highlighted in pink.

**Figure S13. Turnover rate and half-life of the proteins with or without an overlapping SNP-QTL.**

(A, B) The turnover rates (A) and protein half-life values (B) were obtained from Muenzner et al., 2022. The difference was tested using a Wilcoxon test (respective p-values = 0.028 and 0.026).

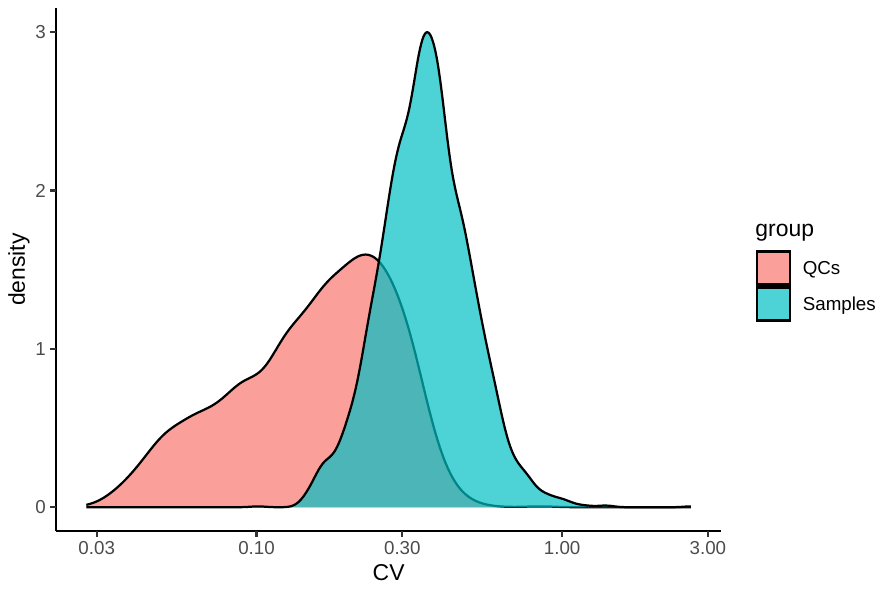

**Figure S14. CV from the QCs and samples precursors.**

The CV was computing either the QCs set or the sample set.
